# *Drosophila* appear resistant to trans-synaptic tau propagation

**DOI:** 10.1101/2024.03.11.584446

**Authors:** James H Catterson, Edmond N Mouofo, Inés López De Toledo Soler, Gillian Lean, Stella Dlamini, Phoebe Liddell, Graham Voong, Taxiarchis Katsinelos, Yu-Chun Wang, Nils Schoovaerts, Patrik Verstreken, Tara L Spires-Jones, Claire S Durrant

## Abstract

Alzheimer’s disease (AD) is the most common cause of dementia in the elderly, prompting extensive efforts to pinpoint novel therapeutic targets for effective intervention. Among the hallmark features of AD is the development of neurofibrillary tangles comprised of hyperphosphorylated tau protein, whose progressive spread throughout the brain is associated with neuronal death. Trans-synaptic propagation of tau has been observed in mouse models and indirect evidence for tau spread via synapses has been observed in human AD. Halting tau propagation is a promising therapeutic target for AD, thus a scalable model system to screen for modifiers of tau spread would be very useful for the field. To this end, we sought to emulate the trans-synaptic spread of human tau (hTau) in *Drosophila melanogaster*. Employing the trans-Tango circuit mapping technique, we investigated whether tau spreads between synaptically connected neurons. Immunohistochemistry and confocal imaging were used to look for tau propagation. Examination of hundreds of flies expressing 4 different human tau constructs in two distinct neuronal populations reveal a robust resistance in *Drosophila* to the trans-synaptic spread of hTau. This resistance persisted in lines with concurrent expression of amyloid-β, in lines with global hTau knock-in to provide a template for human tau in downstream neurons, and with manipulations of temperature. These negative data are important for the field as we establish that *Drosophila* expressing human tau in subsets of neurons are unlikely to be useful to perform screens to find mechanisms to reduce the trans-synaptic spread of tau. The inherent resistance observed in *Drosophila* may serve as a valuable clue, offering insights into strategies for impeding tau spread in future studies.

## Introduction

Accumulation of tau pathology in Alzheimer’s disease (AD) and other tauopathies is associated with neuron death, synapse loss and decline in brain function^1^. In AD, pathological tau appears early in the disease process in the entorhinal cortex and brainstem then progresses to synaptically connected brain regions in such a stereotypical fashion that the pattern of tau pathology is used to determine the stage of disease in post-mortem tissue^2^. This progressive tau spread has also been confirmed longitudinally in living people with AD through positron emission tomography with tracers that bind tau pathology^3–5^. Similarly, in primary tauopathies, tau pathology generally starts in restricted brain regions and spreads through the brain. For example in Progressive Supranuclear Palsy (PSP), post-mortem investigations indicate that tau pathology accumulates early in brainstem, midbrain and basal ganglia and spreads to cerebellum and neocortex later in the disease^6^.

There is converging evidence suggesting that pathological tau spreads between brain regions via synapses by neurons from one brain region releasing tau from pre-synapses which is then taken up by post-synapses in neurons in the downstream brain region. In mice, expression of human tau (hTau) restricted to entorhinal cortex, either by viral injection or use of the neuropsin promotor to drive a transgene, causes both tau accumulation in the entorhinal cortex and the accumulation of tau in downstream brain regions in the hippocampus^7–12^. In mice expressing both hTau and presynaptic vesicles tagged with green fluorescent protein (GFP), we detected hTau in post-synapses in dentate gyrus opposed to hTau-expressing pre-synapses from entorhinal cortex, directly demonstrating trans-synaptic tau spread in this model^11^. Injecting tau derived from post-mortem brain tissue from people who died with tauopathies also induces tau aggregation in mouse models, which spreads from the injection site to connected brain regions^13–16^. While these mouse systems have confounds which could aberrantly drive trans-synaptic spread, there is also mounting human evidence supporting this mechanism of tau propagation. In AD and primary tauopathies PSP and cortico-basal degeneration, MRI studies indicate that tau accumulates in functionally connected brain regions^17–19^. Using high-resolution array tomography imaging, we observe phosphorylated tau within synapses^11,20^, and even find pathological tau within synaptic pairs in brain regions affected late in the disease^21,22^, supporting trans-synaptic spread of tau.

Wherever tau pathology appears in the brain, neurons and synapses die, making preventing tau spread a promising therapeutic target. While mouse model systems robustly exhibit trans-synaptic tau spread, they can take many months to develop a phenotype and are costly and time-consuming models to use for identification of modifiers of tau spread. Human induced pluripotent stem cell derived neurons have also been used to model tau spread^23^, providing a human-relevant *in vitro* model. These cells, however, express largely embryonic forms of tau^24^, which may not be representative of adult physiological functions^25^, and lack intact brain circuits, limiting their utility. *Drosophila melanogaster* has the advantages of short lifespan and a range of molecular tools developed to manipulate gene expression in various cell types. *Drosophila* have been used to study the effects of pathological tau to great effect. Human tau expression in *Drosophila* has been shown in many studies with different tau transgenes and drivers to induce neurodegeneration, and has been used to great utility in identifying modifiers of tau toxicity^26–30^. Of particular interest, tau in *Drosophila* has been implicated in neurodegeneration through binding presynaptic vesicle proteins synaptogyrin-3 and bassoon^31–33^ and synaptic pathways modulate tau toxicity in flies^30^. Tau expressed in the developing *Drosophila* retina has been observed to spread out of the retina into the optic lobe^34,35^, however it was unclear in these studies whether this was trans-synaptic propagation between neurons or whether the observed signal was due to degenerating axon terminals from the retina or glia taking up secreted tau.

In this study, we have harnessed the trans-Tango system in *Drosophila*^36^ to express hTau and GFP in restricted neuronal populations, and tdTomato in synaptically connected downstream neurons, to allow us to address whether tau spreads trans-synaptically in this model system. We further test the influence of several factors that modulate tau spread in mammals in this *Drosophila* system. Amyloid-β (Aβ) pathology has been shown to exacerbate trans-synaptic tau spread in mouse models^37^ and recent data in humans support that synaptic effects of Aβ induce tau accumulation^38^, thus we examine potential tau spread in the context of concurrent human Aβ expression. Although in mice the knockout of mouse tau did not exacerbate human tau spreading phenotypes^39^, we hypothesised that detection of tau spreading in flies could be enhanced if low levels of human tau were present in downstream neurons to allow templated misfolding of the tau spreading from overexpressing neurons. In mice, older animals exhibit more trans-synaptic tau spread^10^ and tau propagation out of the retina was exacerbated by age in *Drosophila*^34^, thus we also examined the effect of ageing in our novel system by examining flies up to 4-6 weeks of age.

## Materials and Methods

### Fly husbandry and stocks

Unless otherwise stated, flies were maintained and all experiments were conducted at 25 °C on a 12 h:12 h light/dark circadian cycle using standard sugar/yeast/agar medium^40^. Only female flies were analysed for the current study. Unless otherwise stated, ‘control’ genotype groups are conditions that use a ‘without hTau’ control cross (typically w^Dah^ or UAS-mCD8:GFP).

The following stocks were obtained from the Bloomington *Drosophila* Stock Center: PDF-GAL4 (#6900, now #80939), Orco-GAL4 (#26818), GH146-QF, QUAS-mCD8:RFP (#30037), UAS-mCD8:GFP (#5137), UAS-mCD8:RFP (#27392), UAS-myr:GFP, QUAS-mtdTom:HA, trans-Tango (#77124), QUAS-Aβ_40_ (#83346), QUAS-Aβ_42_ (#83346), UAS-hTau^WT(0N4R)^ (#78847), UAS-hTau^WT(2N4R)^ (#78861). The control w^Dah^ was a kind gift from Dr Linda Partridge (UCL, UK)^41^. UAS-hTau^P301L(2N4R)^ was a kind gift from Dr Bess Frost (UT Health San Antonio, USA)^42^.

### Generation of hTau transgenic flies

#### Generation of UAS-hTau^0N4R(WT)^:HA and UAS-hTau^0N4R(E14)^:HA flies

For generation of the C-terminally 3xHA-tagged tau transgenic flies, the coding sequence of human tau isoform 0N4R was subcloned into the pJFRC7 vector (Addgene, plasmid #26220. http://n2t.net/addgene:26220; RRID:Addgene_26220, a gift from Gerald Rubin^43^) to allow expression under the control of the GAL4/UAS system. As well as the wild-type version, a phosphomimetic variant of tau (E14) was generated, where 14 serine and threonine residues were mutated to glutamate residues to mimic hyperphosphorylation^44^. The transgenic flies were generated by phiC31 integrase-mediated transgenesis^45^ using attP landing sites 25C6 on the second chromosome and 68A4 on the third chromosome. Plasmid injections and generation of the 3xHA-tagged human tau fly lines were performed by the Cambridge Department of Genetics Fly Facility.

#### Generation of hTau knock-in flies

Human Tau knock-in flies were generated using the CRISPR/Cas9 technique employing a two-step method. Initially, a donor plasmid (pWhite-star) and a tandem gRNA expressing plasmid (pU6-gRNA) were utilized to generate a *Drosophila* Tau (dTau) knock-out. The first exon of dTau was replaced with a mini-white expressing cassette through homology-directed repair (HDR). Subsequently, point mutations of hTau cDNA were introduced using PhiC31-integrase mediated cassette exchange with pKI plasmids.

#### Generation of vectors for *Drosophila* Tau knock-out

Two distinct gRNA sequences targeting dTau were identified:

- gRNA1_Tau: TCAAACGTATGGTCTCTGCA
- gRNA2_Tau: CACTTTTACTTACTAAATTC

These gRNA sequences were introduced into the pU6-BbsI-chiRNA plasmid by restriction/ligation into the BbsI site. The resulting gRNA-expressing plasmid was named pU6-gRNA-Tau. pU6-BbsI-chiRNA was obtained from Addgene, plasmid #45946; http://n2t.net/addgene:45946; RRID:Addgene_45946, a gift from Melissa Harrison, Kate O’Connor-Giles, and Jill Wildonger^46^.

Homology arms of 1 kB surrounding the first exon of dTau were PCR amplified from *Drosophila* genomic DNA using the following primers:

- F_HA1_Tau: GCCACTAGTAGAGACCATACGTTTGACATTCGGG
- R_HA1_Tau: CAGCTCGAGGGCGTTCGGTTCGGGTGT
- F_HA2_Tau: CAGCTCGAGGGCGTTCGGTTCGGGTGT
- R_HA2_Tau: CAGGGTACCGCTACAGCAGCGGAGCATTG

This PCR generated two DNA fragments with SpeI-XhoI and KpnI-AvrII restriction sites, which were used for cloning the fragments into the pWhite-STAR plasmid^47^ via restriction/ligation. The resulting plasmid was named pWhite-STAR_HA-Tau.

#### Generation of vectors for human Tau knock-in

To create the hTau knock-in plasmids, a new backbone containing the 3’ and 5’ UTRs of dTau, with a XhoI restriction site in between for integration of hTau cDNA, was generated. Two fragments were PCR amplified from *Drosophila* genomic DNA using the following primers:

- F_UTR_frag1: CTCACCCATCTGGTCCATCATGATGCTAAGTGCAACAACGCCGAGA
- R_UTR_frag1: CAGCTCGACATGCATTTCGGCATCTCGAGATTGAAAGTCGA
- F_UTR_frag2: GCATTTCGGCATCTCGAGATTGAAAGTCGAACGAGTGTGTGTG
- R_UTR_frag2: CTCACCCATCTGGTCCATCATGATGGCACGGTGATTGCGTCTTG

These fragments were cloned into the MiMIC 1322 pBS-KS-attB1-2 plasmid^48^ after digestion with XbaI and EcoRI, using Gibson assembly. The resulting plasmid was named pKI_UTR_Tau. Next, hTau^WT(0N4R)^ and hTau^WT(2N4R)^ cDNAs were PCR amplified with the following primers:

- F_cDNA_Tau: GACCTCGAGATGGCTGAGCCCCGCCAG
- R_cDNA_Tau: GACCTCGAGTCACAAACCCTGCTTGGCCAG

These PCR products were then cloned into the XhoI restriction site of the pKI_UTR_Tau plasmid via restriction/ligation, resulting in the plasmids pKI_Tau^WT(0N4R)^ and pKI_Tau^WT(2N4R)^. All plasmids were sequence verified, and knock-out and knock-in plasmid injections were performed by BestGene (BestGene Inc., Chino Hills, CA, US).

### Brain dissections and immunostaining

Using forceps, fly heads were removed from the bodies in cold phosphate buffered saline (PBS) containing 4% paraformaldehyde (Pierce, ThermoFisher) + 0.01% Triton X-100 (Sigma). The proboscis was then removed from the fly head to allow fixative and detergent to fix and permeabilize the fly brain. Fly heads were transferred to a 0.6 mL microcentrifuge tube (Fisherbrand™ Premium Microcentrifuge Tubes), fixed in 400 μL 4% paraformaldehyde (PFA) + 0.01% Triton X-100 and rocked for 16 minutes at room temperature. Heads were washed with PBST-1 (PBS containing 0.01% Triton X-100) for 3 × 2 minutes at room temperature. Brains were dissected from the fly heads in ice-cold PBST-1 in a dissection dish under a light stereomicroscope. Brains were transferred to a new 0.6 mL microcentrifuge tube, fixed in 400 μL 4% PFA + 0.1% Triton X-100, and rocked for 20 minutes at room temperature. Brains were then washed in PBST-2 (PBS containing 0.1% Triton X-100) for 3 × 2 minutes and blocked with SeaBlock blocking buffer (ThermoFisher) for 15 minutes at room temperature. SeaBlock blocking buffer was removed and brains were incubated in primary antibody in PBST-2 overnight, washed in PBST-2 for 3 × 20 minutes and incubated in secondary antibody in PBST-2 for 1 hour in the dark. Brains were washed in PBST-2 for 3 × 20 minutes at room temperature in the dark and mounted on a microscope slide using Vectashield mounting media (Vector Labs). To maintain the 3-dimensional structure of the brain, a bridge was made during mounting consisting of two round cover slips (VWR, 13 mm, Thickness 0) which meant the brains were not crushed.

The following antibodies were used. Primary antibodies: Mouse anti-Alz50 (1:250, #Alz50, Courtesy of Dr Peter Davies, Albert Einstein College of Medicine, USA), Mouse anti-AT180 (1:250, #MN1040, ThermoFisher), Mouse anti-AT8 (1:250, #MN1020, ThermoFisher), Chicken anti-GFP (1:1000, #GFP-1020, Aves Labs), Rat anti-HA (1:1000, #11867423001, Roche), Rat anti-NCAD (1:20, #DN-Ex8, Developmental Studies Hybridoma Bank (DSHB)), Rabbit anti-RFP (1:250, # ab62341, Abcam), Rabbit anti-T22 (1:250, Courtesy of Dr Rakez Kayed, University of Texas Medical Branch, USA), Goat anti-Tau (aka Tau Goat, 1:250, #AF3494), Mouse anti-Tau13 (1:250, #835201, BioLegend), Mouse anti-Tau 5A6 (1:50, #5A6, DSHB). Secondary antibodies (all 1:400): Donkey anti-Chicken 488 IgG (#A78948, ThermoFisher), Donkey anti-Goat IgG 647 (#A21447, ThermoFisher), Donkey anti-Mouse 488 IgG (#A21202, ThermoFisher), Donkey anti-Mouse 647 IgM (#ab150123, Abcam), Donkey anti-Mouse 647 IgG (#ab150107, Abcam), Goat anti-Rabbit 546 IgG (#A11035, ThermoFisher), Donkey anti-Rabbit 647 IgG (#A32795, ThermoFisher), Donkey anti-Rat 488 IgG (#A21208, ThermoFisher), Donkey anti-Rat 555 IgG (#A21434, ThermoFisher), Donkey anti-Rat 647 IgG (#A21247, ThermoFisher).

### Microscopy and image analysis

Images were captured with a Leica TCS SP8 confocal microscope (Leica, Wetzlar, Germany) with a 63× oil immersion objective. Images were taken as stacks and are shown as single sections or maximum intensity projections of the stack. All images for one experiment were taken at the same settings. All images were analysed and processed using ImageJ (https://imagej.net/). Brain imaging was performed one brain per genotype, minimizing potential order effects within each genotype. This ensured that all brains within a specific genotype were treated and measured in the same order, preventing any systematic bias related to the order of processing.

We used the ‘Blind Analysis Tools’ ImageJ plug-in (https://imagej.net/plugins/blind-analysis-tools) to blind files before scoring or quantification. Images were ‘Yes/No’ scored for the presence of hTau outside expressing neurons, or for the presence of hTau inside post-synaptic neurons, or for the presence of AT8 in specific subsets of neurons. To measure brain area, we used the ImageJ polygon tool to outline and then measure the part of the brain with the largest area (typically the last slice of the confocal stack). For fluorescence intensity, images were separated into individual channels, thresholded, and then the relevant channel was stacked using Z-stacks with the ‘Sum Slices’ function and then measured to get mean and maximum intensity values. For signal colocalization, we thresholded and segmented individual channels then used the ‘Image Calculator’ function to subtract the PDF signal from the hTau signal. This resulting image was multiplied by the tdTom:HA signal resulting in a new channel representing hTau signal outside PDF neurons and inside post-synaptic neurons. This was then stacked using Z-stacks with the ‘Sum Slices’ function and then measured to get mean and maximum intensity values.

### Study design

Sample sizes for this study were determined by practical limitations and established practice. The number of brains that could be dissected in a single morning was a primary consideration when taking into account potential circadian effects. Sample sizes align with the standard sample size typically used in similar studies within this field. Each data point represents an individual fly brain and no data points were excluded. Each figure legend that contains quantified data has all of the n numbers listed. Randomization was not employed in allocating experimental units to control and treatment groups. While acknowledging the importance of randomization for reducing bias, it was not feasible in this specific study due to the logistical difficulty of randomising fly brains on microscope slides. However, to mitigate potential bias, we implemented blinding during the experiments. For the experiment in Figure 2, microscope slides were blinded by CSD before JHC conducted imaging. In all other instances, the researcher was not fully blinded from the group allocation throughout the experiment. However, as previously mentioned blinding was performed on all acquired images to be analysed or scored.

### Data presentation and statistical analysis

Analysis and visualization of the data was facilitated using the ‘ggplot2’ package in R (RRID:SCR_001905) and RStudio (RRID:SCR_000432, https://posit.co/). The large language models ChatGPT (OpenAI) and Gemini (Google) were employed to assist in refining R code. Data were grouped for each genotype and was displayed as boxplots or stacked bar charts using the ‘ggplot2’, ‘ggbeeswarm’, and ‘ggpattern’ packages. Cartoons were generated using BioRender (RRID:SCR_018361, http://biorender.com). Inkscape (RRID:SCR_014479, http://www.inkscape.org/) was used to generate figures.

Statistical analysis was performed using R and RStudio. Statistical tests used include one-way ANOVA, Kruskal-Wallis rank sum test, Wilcoxon rank sum test, Chi-squared (χ^2^), and Fisher’s exact test, and the appropriate post-hoc analyses were performed. If the model did not meet the assumptions (assessed by plotting histograms to check for normality, plotting model residuals against predictors, normality of residuals was checked with a QQ-plot, and homogeneity of variance checked by plotting residuals against fitted values), data were transformed using the method that was best transformed each individual model to fit the assumptions, for example, Tukey transformation. Statistical details for each individual analyses can be found in the results text. Significance values were reported as p < 0.001 ***.

## Results

### Wildtype and phosphomimetic hTau stays restricted to expressing neurons, even in flies kept at 29°C for 4 weeks

We chose to overexpress hTau in membrane-bound-fluorophore-labelled PDF (Pigment dispersal factor) neurons using the binary GAL4-UAS system and then looked to see if we could observe hTau outside of the expressing neurons. We chose PDF neurons because they are a distinct, well characterised set of neurons (∼16 total)^49^ that also have axon termini located distally from their cell bodies. We focussed on the dorsal axon termini adjacent to the mushroom body calyx as this region is abundant with PDF neuron pre-synapses^50^. In our initial attempts to characterise hTau spread in *Drosophila* we decided to use the monoclonal hTau antibody commonly used in mammalian tau spread studies^37,51,52^ – Tau13 (**Supplementary** Fig. 1A). However, we observed non-specific labelling of structures in post-synaptic neurons and in the mushroom body calyx in controls without hTau overexpression that were not present in no-primary antibody control conditions (**Supplementary** Fig. 1B). We observed a similar non-specificity with the Tau Goat antibody with this antibody detecting apparent cell bodies (**Supplementary** Fig. 1C-D).

**Supplementary Figure 1.**
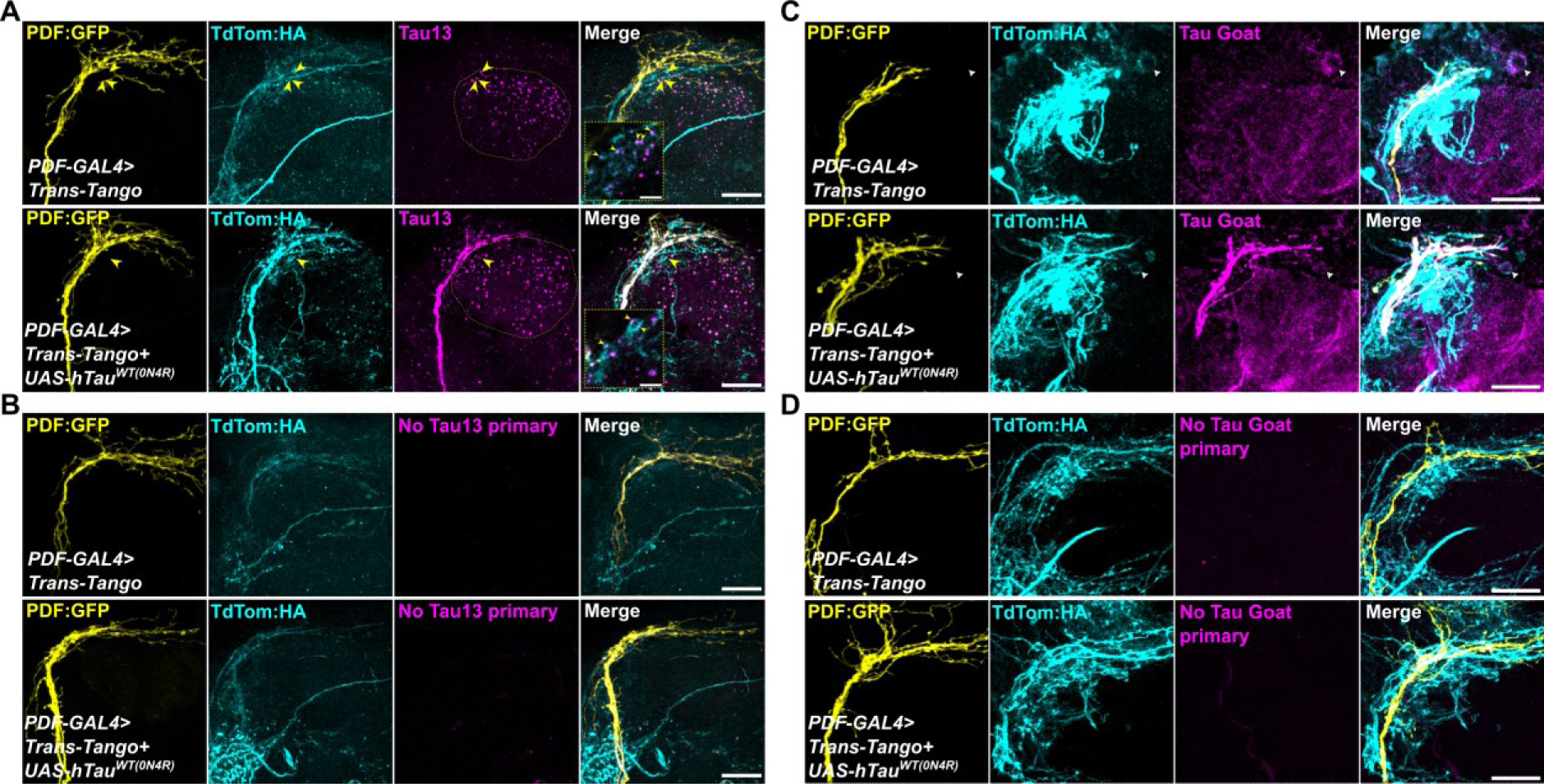
Non-specific binding by human tau antibodies – Tau13 and Tau Goat – in Drosophila brains. (**A**) The hTau-specific antibody, Tau13, labels specific punctate structures in the calyx of the mushroom body (neuropil outlined with dashed yellow in the Tau13 channel). This staining is observed even in conditions that do not contain hTau expression. Some of these puncta colocalise with post-synaptic dendrites (yellow arrowheads). (**B**) Punctate neuropil staining is absent in conditions incubated without the primary antibody (Tau13). (**C**) Tau Goat labels apparent cell bodies in the Kenyon cell area of the mushroom body. Some of these cell bodies colocalise with post-synaptic cell body staining (white arrowheads). This staining is observed even in conditions that do not contain hTau expression. (**D**) Cell body staining is absent in conditions incubated without the primary antibody (Tau Goat). Scale bar is 20 μm for all micrographs except zoomed insets which is 5μm. Genotypes: PDF-GAL4>trans-Tango, PDF-GAL4>trans-Tango + UAS-hTau^WT(0N4R)^.

Thus, we tested other commonly used hTau antibodies (Tau 5A6, AT8, AT180, T22, Alz50) in flies expressing two copies of C-terminal 3xHA-tagged hTau and found that all could label hTau in PDF neurons, though the signal for T22 and Alz50 was reduced likely due to lower levels of oligomeric or misfolded hTau species compared to monomeric forms (**Supplementary** Fig. 2). We decided to use the monoclonal Tau 5A6 antibody to label hTau as it appeared specific and could detect full-length hTau. We confirmed hTau expression in PDF neurons and observed hTau was highly restricted to PDF neurons with no obvious tau spread outside the membrane-GFP signal (**Fig. 1A**). Since we did not observe hTau outside of expressing neurons we hypothesised that increasing temperature could force tau propagation as raising hTau-expressing flies at higher temperatures is known to exacerbate tau toxicity^53^. We generated flies that expressed UAS-mCD8:RFP (to label pre-synaptic neurons) and wildtype UAS-hTau^WT(0N4R)^ or toxic, phosphomimetic UAS-hTau^E14(0N4R)^ (both C-terminally 3xHA-tagged) in PDF neurons and aged flies for 4 weeks. To control for multiple UAS lines, we used a control that expressed both UAS-mCD8:GFP and UAS-mCD8:RFP (**Fig. 1B**). We quantified fluorescence intensity of hTau in each genotype and found an expected significant hTau overexpression in PDF neurons compared to GFP controls (Kruskal-Wallis rank sum test, χ^2^ = 25.2, df = 2, p < 0.0001)(**Fig. 1C**). However, hTau staining outside RFP-labelled PDF neurons was not observed (Chi-square test χ^2^ = 0.063, df = 2, p = 0.97)(**Fig. 1D**).

**Supplementary Figure 2.**
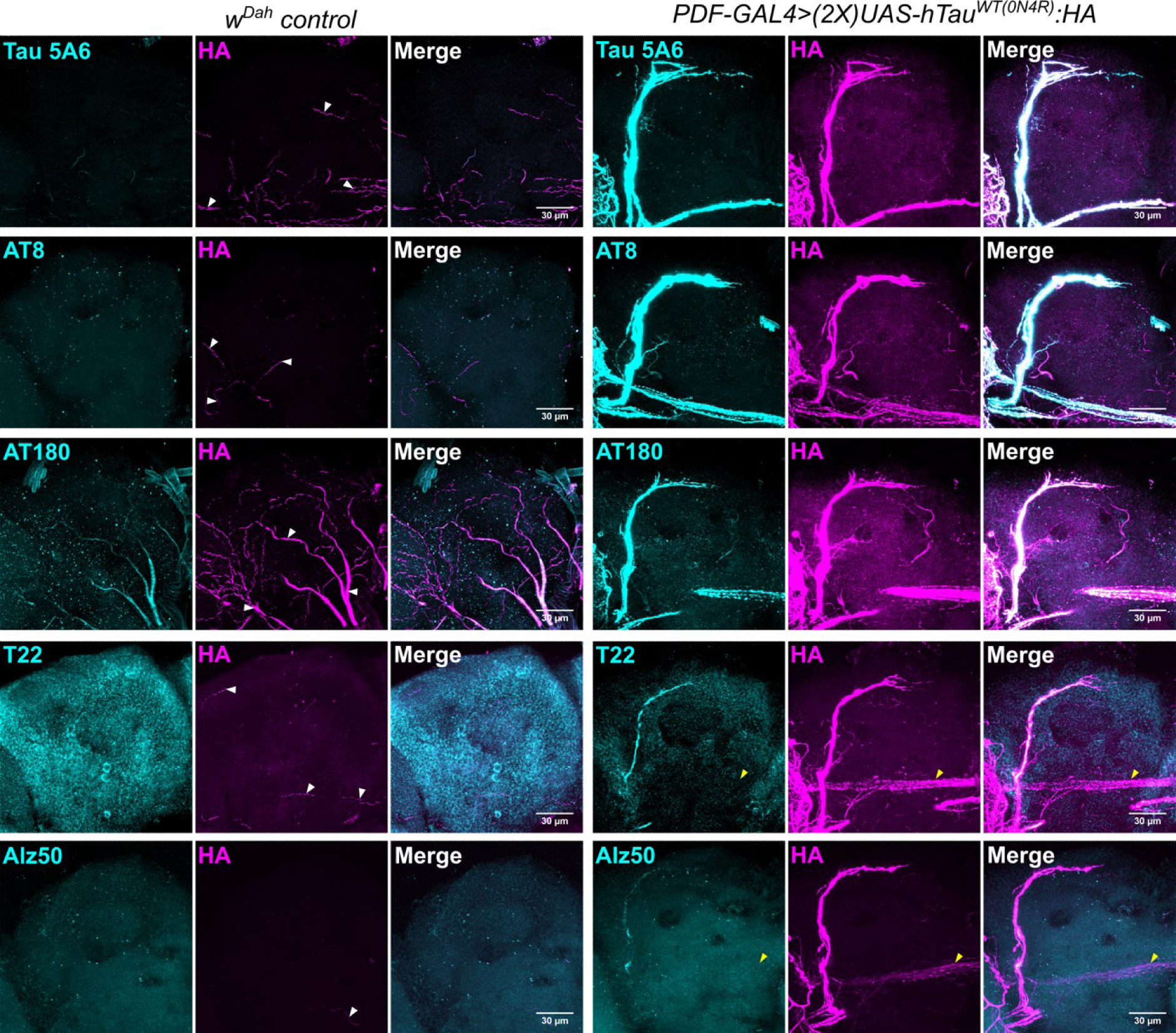
Absence of hTau propagation using different hTau antibodies. Other hTau antibodies could detect HA-tagged hTau expressed in PDF neurons, though the oligomeric (T22) and misfolded (Alz50) forms of hTau had weaker expression and did not fully label all hTau-expressing neurons indicating lower levels. Yellow arrowheads indicate the PDF-expressing posterior optic tract that does not contain staining for oligomeric (T22) or misfolded (Alz50) hTau. Scale bar = 30 μm. White arrowheads indicate autofluorescence from trachea – particularly extensive in the AT180-stained control brain. Genotypes: w^Dah^, PDF-GAL4>UAS-hTau^WT(0N4R)^:HA.

**Figure 1.**
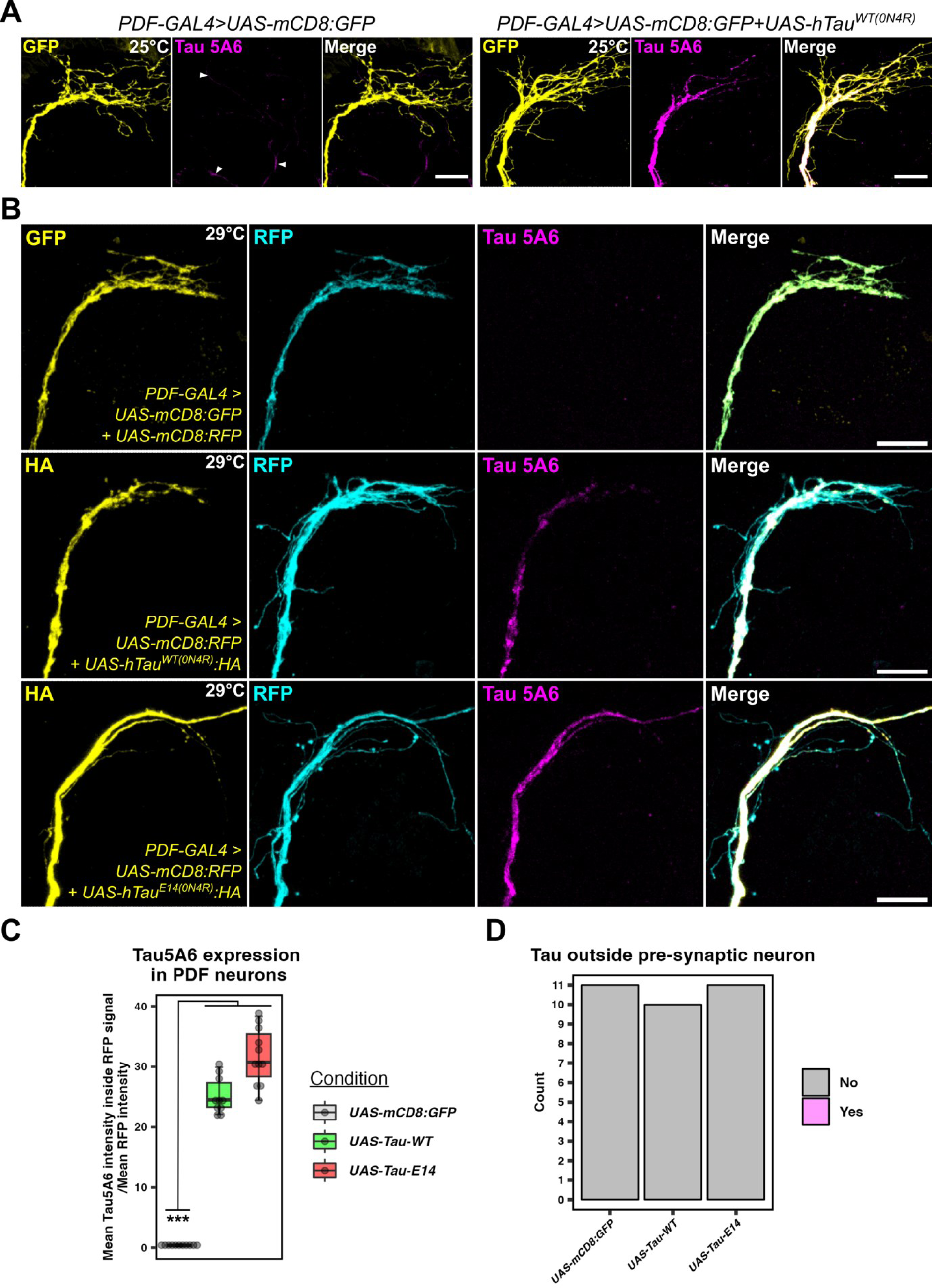
Human Tau stays restricted to expressing neurons, even in flies kept at 29°C for 4 weeks. (**A**) hTau staining was restricted to within the PDF neuron membrane – marked by mCD8:GFP. Control flies without hTau exhibited no Tau 5A6 staining, unlike Tau13-stained controls in **Fig. S1**. (**B**) In flies kept at 29°C for 4 weeks Tau 5A6 staining remained restricted to within the PDF neuron membrane – marked by mCD8:RFP. Control flies without hTau that expressed both GFP and RFP (to control for two UAS lines) exhibited no Tau 5A6 staining. HA was also imaged and was found to overlap with RFP. (**C**) Fluorescence intensity of colocalised Tau 5A6-RFP was measured and normalised to the mean RFP intensity of each brain. Tau 5A6 detected significant hTau overexpression in PDF neurons compared to GFP controls (Kruskal-Wallis rank sum test, χ^2^ = 25.2, df = 2, p < 0.0001; N=10-11 brains). (**D**) Tau 5A6 staining outside RFP-labelled PDF neurons was not observed. Images were blinded, scored and no difference between genotypes was observed (Chi-square test χ^2^ = 0.063, df = 2, p = 0.97; N=10-11 brains). Scale bar is 20 μm. Genotypes: (A) PDF-GAL4>UAS-mCD8:GFP, PDF-GAL4>UAS-mCD8:GFP+UAS-hTau^WT(0N4R)^. (B) PDF-GAL4>UAS-mCD8:GFP+UAS-mCD8:RFP, PDF-GAL4>UAS-mCD8:RFP+UAS-hTau^WT(0N4R)^:HA, PDF-GAL4>UAS-mCD8:RFP+UAS-hTau^E14(0N4R)^:HA.

### Absence of hTau propagation to post-synaptic neurons with different hTau isoforms and different post-synaptic Aβ isoforms

Since we could not detect hTau spreading outside of expressing neurons we hypothesised that a second insult may need to occur to facilitate trans-synaptic hTau spread. We took advantage of the anterograde trans-synaptic tracing tool trans-Tango^36^ to monitor the potential spread of ectopically expressed hTau in PDF neurons (**Fig. 2A-B**). The trans-Tango system labels pre– and post-synaptic neurons with different membrane-tagged fluorophores – GFP and tdTom, respectively. We leveraged the ability of the trans-Tango system’s use of QF expression in post-synaptic neurons to express either QUAS-Aβ_40_ or QUAS-Aβ_42_ as well as QUAS-tdTom (**Fig. 2A**), while expressing UAS-mCD8:GFP and either UAS-hTau^WT(0N4R)^ or UAS-hTau^(2N4R)^ isoforms in pre-synaptic neurons (representative example image in **Fig. 2B**). We quantified total hTau fluorescence and found a significant increase among all hTau-expressing brains regardless of hTau isoform compared to controls without hTau expression (main effect of hTau isoform: one-way ANOVA on Tukey-transformed data: F_2,77_ = 58.73, p < 0.0001; N = 7-10)(**Fig. 2C**). Next, we quantified the amount of hTau outside of the pre-synaptic GFP neurons and colocalising inside of post-synaptic tdTom neurons. We observed no difference in post-synaptic hTau fluorescence between any of the genotypes with all exhibiting very low colocalization signal (main effect of genotype: one-way ANOVA on Tukey-transformed data: F_8,71_ = 0.78, p = 0.62; N = 7-10)(**Fig. 2D**).

**Figure 2.**
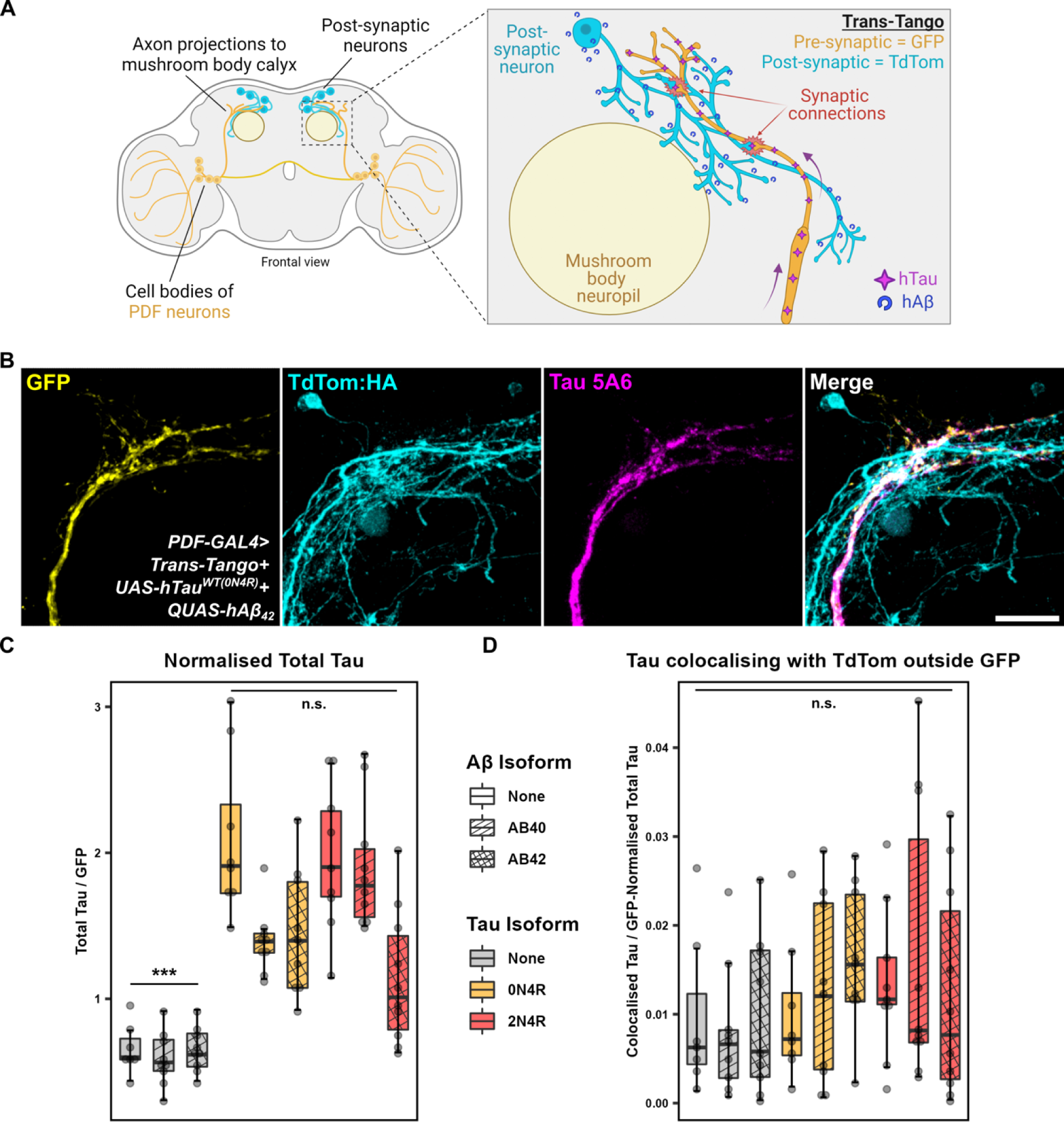
Absence of hTau propagation to post-synaptic neurons with different hTau isoforms and different post-synaptic Aβ isoforms. (**A**) PDF neurons project to the mushroom body calyces. trans-Tango labels pre-synaptic PDF neurons with GFP and their post-synaptic partners with HA-tagged tdTom. Additional experiments using GAL4 and QF drivers (from the trans-Tango system) can also express hTau and hAβ in pre– and post-synaptic compartments. Created using BioRender. (**B**) Representative axon terminal region from PDF neurons highlighting GFP and hTau expression in PDF neurons, and tdTom:HA expression in post-synaptic partners. This fly also expressed Aβ42 in post-synaptic neurons. Staining with Tau 5A6 antibody (full-length hTau) indicated hTau expression in the PDF neuron but not outside. Scale bar = 20 μm. (**C**) Quantification of total hTau fluorescence intensity showed an expected significant increase in hTau-expressing brains compared to controls without hTau (main effect of hTau isoform: one-way ANOVA on Tukey-transformed data: F_2,77_ = 58.73, p < 0.0001; N = 7-10 brains). (**D**) Quantification of hTau colocalization signal outside GFP and inside tdTom signals showed no difference between genotypes (main effect of genotype: one-way ANOVA on Tukey-transformed data: F_8,71_ = 0.78, p = 0.62; N = 7-10 brains). Genotypes (l-r): PDF-GAL4>trans-Tango; PDF-GAL4>trans-Tango+QUAS-Aβ_40_; PDF-GAL4>trans-Tango+QUAS-Aβ_42_; PDF-GAL4>trans-Tango+UAS-hTau^WT(0N4R)^; PDF-GAL4>trans-Tango+UAS-hTau^WT(0N4R)^+QUAS-Aβ_40_; PDF-GAL4>trans-Tango+UAS-hTau^WT(0N4R)^+QUAS-Aβ_42_; PDF-GAL4>trans-Tango+UAS-hTau^WT(2N4R)^; PDF-GAL4>trans-Tango+UAS-hTau^WT(2N4R)^+QUAS-Aβ_40_; PDF-GAL4>trans-Tango+UAS-hTau^WT(2N4R)^+QUAS-Aβ_42_.

### Absence of hTau propagation to post-synaptic neurons in *Drosophila* expressing mutant hTau isoforms combined with global hTau knock-in

It was possible that 16 PDF neurons was too small a population to lead to overt hTau spread so we decided to use the Orco-GAL4 driver to express hTau in ∼1300^54^ olfactory neurons. As with PDF neurons, olfactory neurons also have their cell bodies distal to their axon termini – in this case cell bodies are in the antennae and their axons project to the antennal lobes (**Fig 3A**). We hypothesised that overexpression of hTau could lead to its spread from pre-synaptic olfactory receptor neurons (ORNs) to post-synaptic projection neurons (PNs). Coupled with this, we reasoned that hTau may be required to be present in post-synaptic neurons to be “seeded” in a prion-like manner by trans-synaptic hTau. Thus, we generated Orco-GAL4; hTau^WT(0N4R)^ and Orco-GAL4; hTau^WT(2N4R)^ knock-in recombinant flies that have endogenous *Drosophila* tau (dTau) replaced with hTau (**Fig. 3A**). Two different hTau overexpression transgenes were tested – wildtype UAS-hTau^WT(2N4R)^ and mutant UAS-hTau^P301L(2N4R)^. We stained for AT8 (detects tau phosphorylated at Ser202/Thr205) as we reasoned that staining for total hTau would not be informative due to its abundance in the hTau knock-in. However, we found selective AT8 staining in the hTau^WT(2N4R)^ knock-in controls without hTau overexpression in ORNs (**Supplementary** Fig. 3A). AT8 staining was observed in a small population of neurons dorsolateral to the antennal lobes. We scored the images for the presence of these AT8-stained neurons and found no difference between genotypes (**Supplementary** Fig. 3B, Fisher’s exact, p = 1). Thus, tau phosphorylated at Ser202/Thr205 appeared to naturally accumulate in this neuronal subset, so it was necessary to be able to identify post-synaptic PNs to detect potential hTau spread. To make sure we could correctly identify PNs, we combined Orco-GAL4; hTau^WT^ recombinants with GH146-QF>QUAS-mCD8:RFP recombinants which drive RFP expression in PNs. For this experiment we used the GAL4/UAS and QF/QUAS binary expression systems because the combination of all the required transgenes could not be easily combined with trans-Tango. Our Orco-GAL4/GH146-QF>QUAS-mCD8:RFP; hTau^WT^ recombinants were crossed with w^Dah^ controls and UAS-hTau^WT(0N4R)^:HA flies. Thus, we aimed to detect trans-synaptic hTau spread from ORNs to RFP-labelled PNs in a global hTau knock-in background. As before, AT8 stained neurons dorsolateral to the antennal lobes, but these were not overlapping with post-synaptic PNs (see dashed yellow areas in **Fig. 3B**). We observed robust AT8 staining in ORNs from flies overexpressing hTau^WT(0N4R)^(Wilcoxon rank sum exact test, W = 0, p = 0.0012, **Fig. 3C**), but AT8 did not capture trans-synaptic hTau spread (Chi-square test χ^2^ = 0.077, df = 1, p = 0.78, **Fig. 3D**).

**Supplementary Figure 3.**
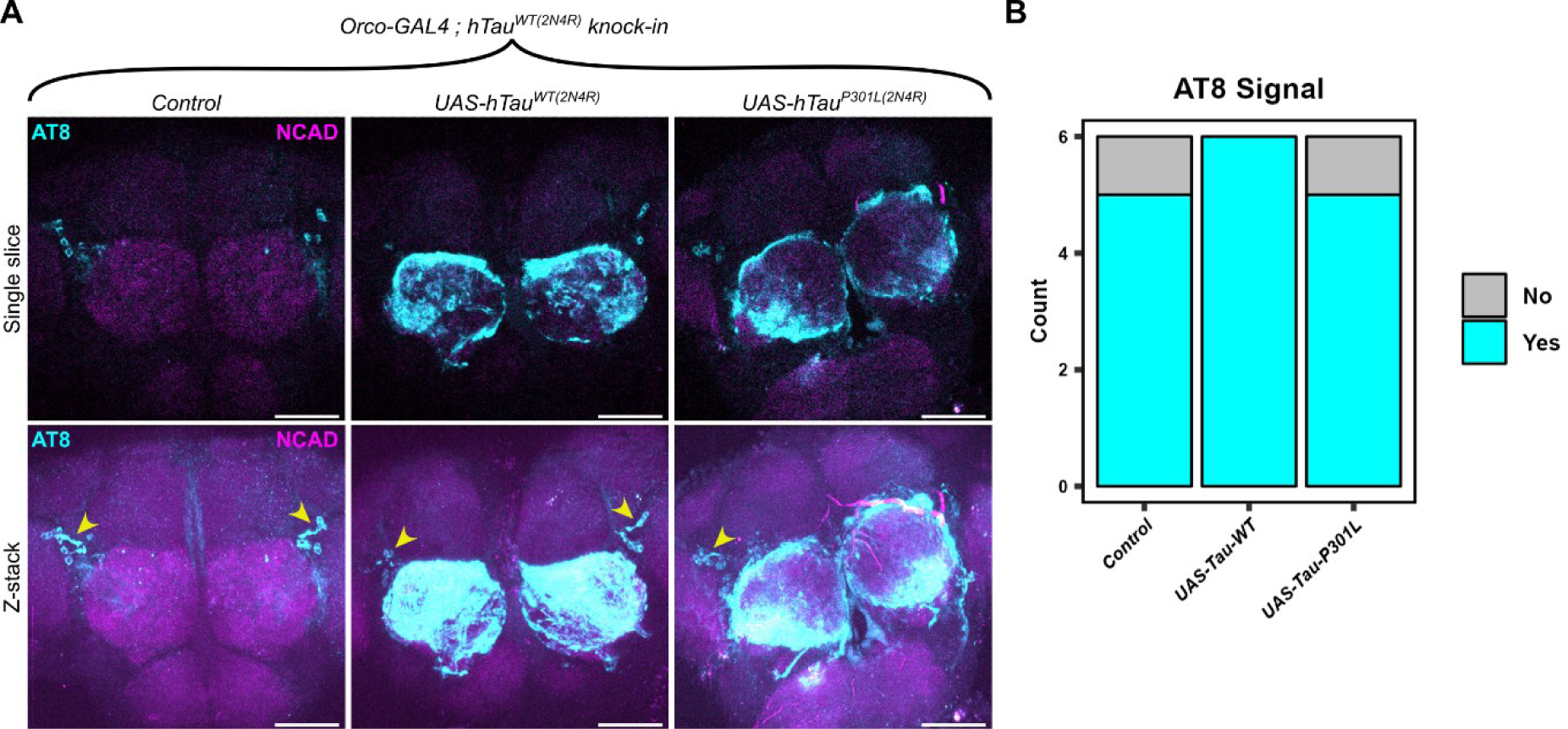
AT8 staining in hTau knock-in flies. (**A**) Single confocal slices and z-stacks from 6-week-old flies heterozygous for hTau^WT(2N4R)^ knock-in. We observed consistent AT8 staining in a small subset of neurons dorsolaterally to the antennal lobes that was present even in controls without hTau overexpression. (**B**) AT8 staining of the region surrounding the antennal lobes was found in all genotypes. Fisher’s exact test, chosen for its suitability with small sample size, detected no significant association between genotype and AT8 signal (p = 1; N=6 brains). Scale bar is 50 μm. Genotypes: Orco-GAL4/+; hTau^WT(2N4R)^, Orco-GAL4>UAS-hTau^WT(2N4R)^; hTau^WT(2N4R)^, Orco-GAL4>UAS-hTau^P301L(2N4R)^; hTau^WT(2N4R)^.

**Figure 3.**
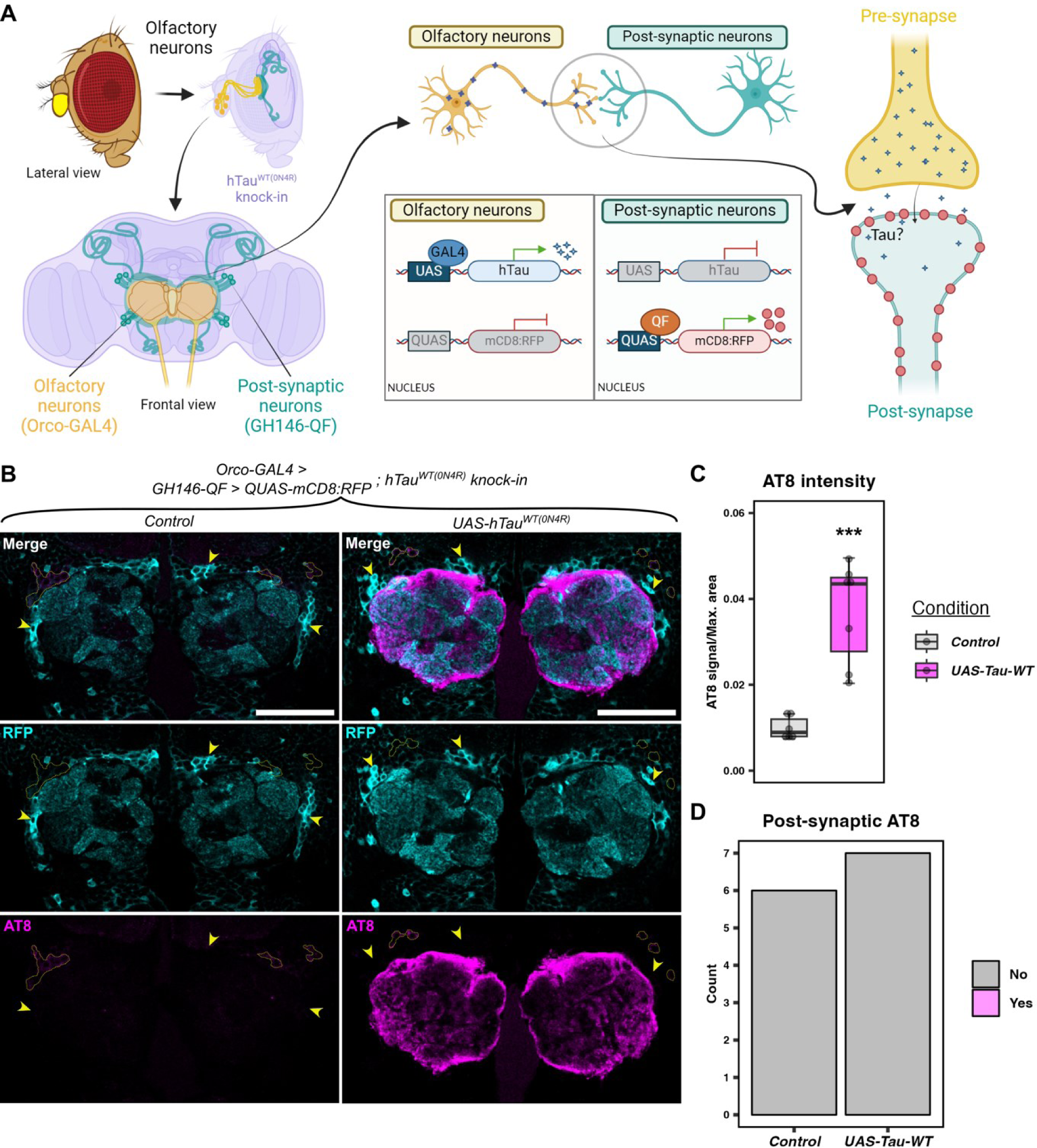
Absence of hTau propagation in post-synaptic neurons. (**A**) Schematic shows the brain regions involved in the experiment. Yellow neurons represent olfactory neurons labelled by Orco-GAL4, while teal neurons represent projection neurons connecting the antennal lobes to the mushroom body, labelled by GH146-QF. The background colour (lilac) indicates the presence of a global hTau^WT(0N4R)^ knock-in mutation. Created using BioRender. (**B**) In a global hTau^WT(0N4R)^ heterozygous background, GH146-QF labels post-synaptic neurons with mCD8:RFP while pre-synaptic Orco-GAL4 expresses hTau^WT(0N4R)^ or control. AT8 did not detectably spread trans-synaptically. Yellow arrowheads and yellow-dashed areas label clusters of post-synaptic cell bodies and endogenous hTau accumulation, respectively. (**C**) Fluorescence intensity of AT8 was measured and normalised to the maximum area of each brain. AT8 detected significant hTau^WT^ overexpression in the antennal lobes compared to control flies (Wilcoxon rank sum exact test, W = 0, p = 0.0012; N=6-7 brains). (**D**) AT8 staining within RFP-labelled post-synaptic neurons was not observed. Images were blinded, scored and no difference between genotypes was observed (Chi-square test χ^2^ = 0.077, df = 1, p = 0.78; N=6-7 brains). Scale bar is 50 μm. Genotypes: Orco-GAL4; GH146-QF>QUAS-mCD8:RFP; hTau^WT(0N4R)^, Orco-GAL4>UAS-hTau^WT(0N4R)^; GH146-QF>QUAS-mCD8:RFP; hTau^WT(0N4R)^, Orco-GAL4>UAS-hTau^E14(0N4R)^; GH146-QF>QUAS-mCD8:RFP; hTau^WT(0N4R)^.

## Discussion

In this work, we sought to generate a *Drosophila* model of trans-synaptic tau spread. Despite testing multiple different experimental paradigms, we found that *Drosophila* are remarkably resistant to the trans-synaptic tau spread evidenced in human tauopathies and observed in mammalian model systems^3–12,16,17,21,22,55,56^. This important negative data demonstrates that, whilst exceptionally useful for a number of other studies in neurodegeneration, including modifiers of tau toxicity^57–60^, *Drosophila* are unlikely to be a suitable model for screening modifiers of trans-synaptic tau spread. In addition, the resistance to spread seen in this model, compared to mammalian systems, opens up future possibilities to determine factors missing from *Drosophila,* such as proteins, cell types or neuronal functionality^61,62^ that could provide insight into mammalian mechanisms. This also contrasts with fly models successfully showing spread and seeding of toxic neuropathological proteins such as huntingtin^63–65^, FUS^66^, TDP-43^67^, and Aβ^68^.

In this study we perform a robust assessment of suitable tau antibodies for immunostaining experiments. Crucially, we found that the commonly used human Tau antibodies, Tau13^69^ and Tau Goat^22^, showed significant non-specific staining, even in *Drosophila* lacking hTau. This suggests that both antibodies recognise additional non-tau epitopes in *Drosophila,* and should be avoided where possible. In this study, we find that using either Tau 5A6 as a full-length tau antibody, or AT8 to detect phosphorylated tau, provided optimal, specific staining.

In our work, we studied the effect of multiple types of hTau to maximise the likelihood of observing trans-synaptic tau spread. When wildtype 0N4R hTau was overexpressed in PDF neurons, we did not find evidence of hTau spreading beyond the confines of PDF neurons in up to 4-week-old flies. Interestingly, despite tau phosphorylation appearing to be an important process in permitting tau spread in human disease and mammalian models^70,71^, we also did not find any evidence of tau spread in flies expressing phosphomimetic (E14) 0N4R hTau. Nor did we find evidence of tau spread in flies expressing mutant P301L 2N4R hTau. To further push the system, we tested the hypothesis that hTau may require an hTau template in the post-synaptic compartment in order to “seed” into the post synapse. Whilst studies in mice have found that hTau can spread without mouse tau present in the post-synaptic compartment^39^, hTau may be required to observe effective tau seeding^55,72^. We replaced dTau with a wildtype hTau knock in in all neuronal cells, but still did not detect movement of phosphorylated tau from the presynaptic neurons into postsynaptic structures. Taken together, we find that, regardless of tau length, phosphorylation status, mutation status or combination with hTau at the post-synapse, *Drosophila* do not show significant trans-synaptic tau spread.

A key advantage of model systems like *Drosophila* is the ability to finely control the experimental environment. Previous studies have shown that raising flies under higher temperatures can exacerbate tau toxicity^53^, so we performed follow-up experiments in this “high stress” environment. Interestingly, increasing the temperature had no impact on the detection of wildtype 0N4R hTau, or E14 0N4R hTau outside of the PDF neurons.

Given studies in mice and humans have indicated that Aβ can enhance levels of tau spread and toxicity^37,38^, we examined whether the addition of Aβ to our *Drosophila* model could be sufficient to promote tau spread. Once again, we found that, regardless of the length of wildtype hTau expressed in PDF neurons (0N4R compared with 2N4R), or the expression of either human Aβ_40_ or Aβ_42_ from postsynaptic compartments, we could not detect hTau outside of the PDF neurons. This suggests that, in *Drosophila*, combining the two key pathogenic hallmarks of AD is not sufficient to induce tau spread, although evidence from other studies show you can still gain useful insights on the impacts of Aβ and tau on neuronal toxicity and synaptic function^59,60^.

The majority of work in this study was performed using the well-characterised PDF neurons as the primary site for overexpressing tau. PDF neurons are a set of ∼16 neurons known to play key roles in maintaining circadian rhythms^49^. PDF neurons were chosen as they have axon termini located distally from their cell bodies, allowing us observe presynaptic structures, and their immediate postsynaptic environment, in isolation from the rest of the cell. This permits for more effective assessment of whether tau signals co-localise within the PDF neuron, or whether exogenous tau signal represents genuine spread to post-synaptic compartments. As we did not see tau spread from PDF neurons, we sought to establish whether this is a quirk of this specific neuronal subtype, or a consequence of it being a small neuronal population. We performed additional experiments expressing hTau in olfactory neurons, comprising of around 1300 individual cells with axons projecting distal to their cell bodies^54^. In our hands, we did not observe phosphorylated tau in postsynaptic projection neurons. Taken together, we show that, in two well characterised neuronal populations in *Drosophila*, overexpression of hTau does not lead to spread of tau to post-synaptic compartments. Whilst we cannot rule out other neuronal circuits being amenable to tau spread, inability to study distal axonal projections will introduce significant noise while imaging, rendering them unsuitable for screening studies. Previous studies have examined the *Drosophila* eye for evidence of tau spread, and have shown that tau deposits appear to move from the retina and lamina into the optic lobe during aging, with the distance migrated reduced when tau phosphorylation is lowered^34^. It is unclear, however, whether the migration of tau over a short distance reported in this paper is a consequence of genuine trans-synaptic spread, or whether tau released from dying neurons has been taken up by other cells.

Overall, we conclude that *Drosophila* are highly resistant to trans-synaptic spread of tau in two major central nervous system neuronal populations. This model, whilst highly useful for studies of toxicity^59,60^, may therefore not be appropriate to conduct high throughput screens looking for modifiers of tau spread. Examining why *Drosophila* do not show tau spread as seen in other model systems, may provide mechanistic insight that could inform therapeutic development.

## Data availability

All relevant data can be found within the article and its supplementary information and will be available in Edinburgh University Data Repository.

## Abbreviations

Aβ: Amyloid beta peptide
AD: Alzheimer’s disease
ANOVA: Analysis of variance
E14: Phosphomimetic mutant tau
GFP: Green fluorescent protein
HA: Human influenza hemagglutinin
IgG: Immunoglobulin G
MRI: Magnetic resonance imaging
ORN: Olfactory receptor neuron
P301L: Pathogenic mutant tau
PBS: Phosphate buffered saline
PBST: Phosphate buffered saline with Triton-X
PDF: Pigment dispersing factor neuropeptide
PFA: Paraformaldehyde
PN: Projection neuron
PSP: Progressive supranuclear palsy
RFP: Red fluorescent protein
WT: Wildtype

## Acknowledgements

We are grateful to past and present members of the Durrant and Spires-Jones labs for helpful discussions. We also thank Michael Molinek and Krisztina Vinko for management and delivery of the *Drosophila* food, and Jane Tulloch for administrative support. Thanks to Dr Thomas R Jahn for his support in the generation of the UAS-hTau^WT(0N4R)^:HA and UAS-hTau^E14(0N4R)^:HA fly lines. We also thank the wider fly community for the generous sharing of reagents and stocks, particularly the University of Cambridge Department of Genetics Fly Facility and the Bloomington *Drosophila* Stock Center (NIH P40OD018537).

## References

1. Tzioras M, McGeachan RI, Durrant CS, Spires-Jones TL. Synaptic degeneration in Alzheimer disease. Nat Rev Neurol. 2023;19(1):19–38. doi:10.1038/s41582-022-00749-z

2. Braak H, Braak E. Neuropathological stageing of Alzheimer-related changes. Acta Neuropathol. 1991;82(4):239–259. doi:10.1007/BF00308809

3. Oh M, Oh SJ, Lee SJ, et al. One-Year Longitudinal Changes in Tau Accumulation on [18F]PI-2620 PET in the Alzheimer Spectrum. Journal of Nuclear Medicine. Published online February 1, 2024. doi:10.2967/jnumed.123.265893

4. Macedo AC, Tissot C, Therriault J, et al. The Use of Tau PET to Stage Alzheimer Disease According to the Braak Staging Framework. J Nucl Med. 2023;64(8):1171–1178. doi:10.2967/jnumed.122.265200

5. Biel D, Brendel M, Rubinski A, et al. Tau-PET and in vivo Braak-staging as prognostic markers of future cognitive decline in cognitively normal to demented individuals. Alzheimer’s Research & Therapy. 2021;13(1):137. doi:10.1186/s13195-021-00880-x

6. Kovacs GG, Lukic MJ, Irwin DJ, et al. Distribution patterns of tau pathology in progressive supranuclear palsy. Acta Neuropathol. 2020;140(2):99–119. doi:10.1007/s00401-020-02158-2

7. Calignon A, Polydoro M, Suárez-Calvet M, et al. Propagation of tau pathology in a model of early Alzheimer’s disease. Neuron. 2012;73:685–697.

8. Liu L, Drouet V, Wu JW, et al. Trans-synaptic spread of tau pathology in vivo. PLoS One. 2012;7:31302.

9. Harris JA, Koyama A, Maeda S, et al. Human P301L-mutant tau expression in mouse entorhinal-hippocampal network causes tau aggregation and presynaptic pathology but no cognitive deficits. PLoS One. 2012;7:45881.

10. Wegmann S, Bennett RE, Delorme L, et al. Experimental evidence for the age dependence of tau protein spread in the brain. Science Advances. 2019;5(6):eaaw6404. doi:10.1126/sciadv.aaw6404

11. Pickett EK, Henstridge CM, Allison E, et al. Spread of tau down neural circuits precedes synapse and neuronal loss in the rTgTauEC mouse model of early Alzheimer’s disease. Published online 2017.

12. Davies C, Tulloch J, Yip E, et al. Apolipoprotein E isoform does not influence trans-synaptic spread of tau pathology in a mouse model. Brain Neurosci Adv. 2023;7:23982128231191050. doi:10.1177/23982128231191046

13. Clavaguera F, Akatsu H, Fraser G, et al. Brain homogenates from human tauopathies induce tau inclusions in mouse brain. Proc Natl Acad Sci U S A. 2013;110(23):9535–9540. doi:10.1073/pnas.1301175110

14. Narasimhan S, Guo JL, Changolkar L, et al. Pathological Tau Strains from Human Brains Recapitulate the Diversity of Tauopathies in Nontransgenic Mouse Brain. J Neurosci. 2017;37:11406–11423.

15. Boluda S, Iba M, Zhang B, Raible KM, Lee VMY, Trojanowski JQ. Differential induction and spread of tau pathology in young PS19 tau transgenic mice following intracerebral injections of pathological tau from Alzheimer’s disease or corticobasal degeneration brains. Acta Neuropathol. 2015;129:221–237.

16. Ahmed Z, Cooper J, Murray TK, et al. A novel in vivo model of tau propagation with rapid and progressive neurofibrillary tangle pathology: the pattern of spread is determined by connectivity, not proximity. Acta Neuropathol. 2014;127:667–683.

17. Franzmeier N, Neitzel J, Rubinski A, et al. Functional brain architecture is associated with the rate of tau accumulation in Alzheimer’s disease. Nat Commun. 2020;11:347.

18. Franzmeier N, Rubinski A, Neitzel J, et al. Functional connectivity associated with tau levels in ageing, Alzheimer’s, and small vessel disease. Brain. 2019;142:1093–1107.

19. Franzmeier N, Brendel M, Beyer L, et al. Tau deposition patterns are associated with functional connectivity in primary tauopathies. Nat Commun. 2022;13(1):1362. doi:10.1038/s41467-022-28896-3

20. Taylor LW, Simzer EM, Pimblett C, et al. p-tau Ser356 is associated with Alzheimer’s disease pathology and is lowered in brain slice cultures using the NUAK inhibitor WZ4003. Acta Neuropathol. 2024;147(1):7. doi:10.1007/s00401-023-02667-w

21. Colom-Cadena M, Davies C, Sirisi S, et al. Synaptic oligomeric tau in Alzheimer’s disease — A potential culprit in the spread of tau pathology through the brain. Neuron. 2023;111(14):2170–2183.e6. doi:10.1016/j.neuron.2023.04.020

22. McGeachan RI, Keavey L, Rose JL, et al. Evidence for trans-synaptic propagation of oligomeric tau in Progressive Supranuclear Palsy. Published online February 12, 2024. doi:10.1101/2022.09.20.22280086

23. Rauch JN, Luna G, Guzman E, et al. LRP1 is a master regulator of tau uptake and spread. Nature. 2020;580:381–385.

24. Sposito T, Preza E, Mahoney CJ, et al. Developmental regulation of tau splicing is disrupted in stem cell-derived neurons from frontotemporal dementia patients with the 10+16 splice-site mutation in MAPT. Hum Mol Genet. 2015;24(18):5260–5269. doi:10.1093/hmg/ddv246

25. Kent SA, Spires-Jones TL, Durrant CS. The physiological roles of tau and Aβ: implications for Alzheimer’s disease pathology and therapeutics. Acta Neuropathol. Published online July 29, 2020. doi:10.1007/s00401-020-02196-w

26. Fulga TA, Elson-Schwab I, Khurana V, et al. Abnormal bundling and accumulation of F-actin mediates tau-induced neuronal degeneration in vivo. Nat Cell Biol. 2007;9(2):139–148. doi:10.1038/ncb1528

27. Wittmann CW, Wszolek MF, Shulman JM, et al. Tauopathy in Drosophila: neurodegeneration without neurofibrillary tangles. Science. 2001;293(5530):711–714. doi:10.1126/science.1062382

28. Butzlaff M, Hannan SB, Karsten P, et al. Impaired retrograde transport by the Dynein/Dynactin complex contributes to Tau-induced toxicity. Hum Mol Genet. 2015;24(13):3623–3637. doi:10.1093/hmg/ddv107

29. Shulman JM, Chipendo P, Chibnik LB, et al. Functional Screening of Alzheimer Pathology Genome-wide Association Signals in Drosophila. Am J Hum Genet. 2011;88(2):232–238. doi:10.1016/j.ajhg.2011.01.006

30. Praschberger R, Kuenen S, Schoovaerts N, et al. Neuronal identity defines α-synuclein and tau toxicity. Neuron. 2023;111(10):1577–1590.e11. doi:10.1016/j.neuron.2023.02.033

31. Martinez P, Patel H, You Y, et al. Bassoon contributes to tau-seed propagation and neurotoxicity. Nat Neurosci. 2022;25(12):1597–1607. doi:10.1038/s41593-022-01191-6

32. Zhou L, McInnes J, Wierda K, et al. Tau association with synaptic vesicles causes presynaptic dysfunction. Nat Commun. 2017;8:15295. doi:10.1038/ncomms15295

33. Largo-Barrientos P, Apóstolo N, Creemers E, et al. Lowering Synaptogyrin-3 expression rescues Tau-induced memory defects and synaptic loss in the presence of microglial activation. Neuron. Published online January 7, 2021. doi:10.1016/j.neuron.2020.12.016

34. Aqsa, Sarkar S. Age dependent trans-cellular propagation of human tau aggregates in Drosophila disease models. Brain Research. 2021;1751:147207. doi:10.1016/j.brainres.2020.147207

35. Kilian JG, Hsu HW, Mata K, Wolf FW, Kitazawa M. Astrocyte transport of glutamate and neuronal activity reciprocally modulate tau pathology in Drosophila. Neuroscience. 2017;348:191–200. doi:10.1016/j.neuroscience.2017.02.011

36. Talay M, Richman EB, Snell NJ, et al. Transsynaptic Mapping of Second-Order Taste Neurons in Flies by trans-Tango. Neuron. 2017;96(4):783–795.e4. doi:10.1016/j.neuron.2017.10.011

37. Pooler AM, Polydoro M, Maury EA, et al. Amyloid accelerates tau propagation and toxicity in a model of early Alzheimer’s disease. Acta Neuropathologica Communications. 2015;3(1):14. doi:10.1186/s40478-015-0199-x

38. Giorgio J, Adams JN, Maass A, Jagust WJ, Breakspear M. Amyloid induced hyperexcitability in default mode network drives medial temporal hyperactivity and early tau accumulation. Neuron. 2024;112(4):676–686.e4. doi:10.1016/j.neuron.2023.11.014

39. Wegmann S, Maury EA, Kirk MJ, et al. Removing endogenous tau does not prevent tau propagation yet reduces its neurotoxicity. EMBO J. 2015;34:3028–3041.

40. Catterson JH, Khericha M, Dyson MC, et al. Short-Term, Intermittent Fasting Induces Long-Lasting Gut Health and TOR-Independent Lifespan Extension. Curr Biol. 2018;28(11):1714–1724.e4. doi:10.1016/j.cub.2018.04.015

41. Catterson JH, Minkley L, Aspe S, et al. Protein retention in the endoplasmic reticulum rescues Aβ toxicity in Drosophila. Neurobiology of Aging. 2023;132:154–174. doi:10.1016/j.neurobiolaging.2023.09.008

42. Levy SA, Zuniga G, Gonzalez EM, Butler D, Temple S, Frost B. TauLUM, an in vivo Drosophila sensor of tau multimerization, identifies neuroprotective interventions in tauopathy. Cell Rep Methods. 2022;2(9):100292. doi:10.1016/j.crmeth.2022.100292

43. Pfeiffer BD, Ngo TTB, Hibbard KL, et al. Refinement of Tools for Targeted Gene Expression in Drosophila. Genetics. 2010;186(2):735–755. doi:10.1534/genetics.110.119917

44. Khurana V, Lu Y, Steinhilb ML, Oldham S, Shulman JM, Feany MB. TOR-Mediated Cell-Cycle Activation Causes Neurodegeneration in a *Drosophila* Tauopathy Model. Current Biology. 2006;16(3):230–241. doi:10.1016/j.cub.2005.12.042

45. Bischof J, Maeda RK, Hediger M, Karch F, Basler K. An optimized transgenesis system for Drosophila using germ-line-specific φC31 integrases. Proceedings of the National Academy of Sciences. 2007;104(9):3312–3317. doi:10.1073/pnas.0611511104

46. Gratz SJ, Cummings AM, Nguyen JN, et al. Genome Engineering of Drosophila with the CRISPR RNA-Guided Cas9 Nuclease. Genetics. 2013;194(4):1029–1035. doi:10.1534/genetics.113.152710

47. Choi CM, Vilain S, Langen M, et al. Conditional Mutagenesis in Drosophila. Science. 2009;324(5923):54–54. doi:10.1126/science.1168275

48. Venken KJT, Schulze KL, Haelterman NA, et al. MiMIC: a highly versatile transposon insertion resource for engineering Drosophila melanogaster genes. Nat Methods. 2011;8(9):737–743. doi:10.1038/nmeth.1662

49. Helfrich-Förster C. Robust circadian rhythmicity of Drosophila melanogaster requires the presence of lateral neurons: a brain-behavioral study of disconnected mutants. J Comp Physiol A. 1998;182(4):435–453. doi:10.1007/s003590050192

50. Shafer OT, Gutierrez GJ, Li K, et al. ---Connectomic analysis of the Drosophila lateral neuron clock cells reveals the synaptic basis of functional pacemaker classes. Desplan C, ed. eLife. 2022;11:e79139. doi:10.7554/eLife.79139

51. Dujardin S, Fernandes A, Bannon R, et al. Tau propagation is dependent on the genetic background of mouse strains. Brain Communications. 2022;4(2):fcac048. doi:10.1093/braincomms/fcac048

52. Takeda S, Wegmann S, Cho H, et al. Neuronal uptake and propagation of a rare phosphorylated high-molecular-weight tau derived from Alzheimer’s disease brain. Nat Commun. 2015;6:8490.

53. Kosmidis S, Grammenoudi S, Papanikolopoulou K, Skoulakis EMC. Differential Effects of Tau on the Integrity and Function of Neurons Essential for Learning in Drosophila. J Neurosci. 2010;30(2):464–477. doi:10.1523/JNEUROSCI.1490-09.2010

54. Vosshall LB. The Molecular Logic of Olfaction in Drosophila. Chemical Senses. 2001;26(2):207–213. doi:10.1093/chemse/26.2.207

55. Miller LVC, Mukadam AS, Durrant CS, et al. Tau assemblies do not behave like independently acting prion-like particles in mouse neural tissue. Acta Neuropathologica Communications. 2021;9(1):41. doi:10.1186/s40478-021-01141-6

56. Mudher A, Colin M, Dujardin S, et al. What is the evidence that tau pathology spreads through prion-like propagation? Acta Neuropathologica Communications. 2017;5(1):99. doi:10.1186/s40478-017-0488-7

57. Feuillette S, Charbonnier C, Frebourg T, Campion D, Lecourtois M. A Connected Network of Interacting Proteins Is Involved in Human-Tau Toxicity in Drosophila. Front Neurosci. 2020;14:68. doi:10.3389/fnins.2020.00068

58. Lasagna-Reeves CA, de Haro M, Hao S, et al. Reduction of Nuak1 decreases tau and reverses phenotypes in a tauopathy mouse model. Neuron. 2016;92(2):407–418. doi:10.1016/j.neuron.2016.09.022

59. Gistelinck M, Lambert JC, Callaerts P, Dermaut B, Dourlen P. *Drosophila* Models of Tauopathies: What Have We Learned? International Journal of Alzheimer’s Disease. 2012;2012:e970980. doi:10.1155/2012/970980

60. Prüßing K, Voigt A, Schulz JB. Drosophila melanogaster as a model organism for Alzheimer’s disease. Molecular Neurodegeneration. 2013;8(1):35. doi:10.1186/1750-1326-8-35

61. Ugur B, Chen K, Bellen HJ. Drosophila tools and assays for the study of human diseases. Disease Models & Mechanisms. 2016;9(3):235–244. doi:10.1242/dmm.023762

62. Freeman MR, Doherty J. Glial cell biology in Drosophila and vertebrates. Trends in Neurosciences. 2006;29(2):82–90. doi:10.1016/j.tins.2005.12.002

63. Donnelly KM, DeLorenzo OR, Zaya AD, et al. Phagocytic glia are obligatory intermediates in transmission of mutant huntingtin aggregates across neuronal synapses. Bellen HJ, Westbrook GL, eds. eLife. 2020;9:e58499. doi:10.7554/eLife.58499

64. Pearce MMP, Spartz EJ, Hong W, Luo L, Kopito RR. Prion-like transmission of neuronal huntingtin aggregates to phagocytic glia in the Drosophila brain. Nat Commun. 2015;6(1):6768. doi:10.1038/ncomms7768

65. Babcock DT, Ganetzky B. Transcellular spreading of huntingtin aggregates in the Drosophila brain. Proceedings of the National Academy of Sciences. 2015;112(39):E5427–E5433. doi:10.1073/pnas.1516217112

66. Feuillette S, Delarue M, Riou G, et al. Neuron-to-Neuron Transfer of FUS in Drosophila Primary Neuronal Culture Is Enhanced by ALS-Associated Mutations. J Mol Neurosci. 2017;62(1):114–122. doi:10.1007/s12031-017-0908-y

67. Chang YH, Dubnau J. Endogenous retroviruses and TDP-43 proteinopathy form a sustaining feedback driving intercellular spread of Drosophila neurodegeneration. Nat Commun. 2023;14(1):966. doi:10.1038/s41467-023-36649-z

68. Sowade RF, Jahn TR. Seed-induced acceleration of amyloid-β mediated neurotoxicity in vivo. Nat Commun. 2017;8(1):512. doi:10.1038/s41467-017-00579-4

69. Michael J. Ellis, Christiana Lekka, Hanna Tulmin, et al. Validation of Tau Antibodies for Use in Western Blotting and Immunohistochemistry. bioRxiv. Published online January 1, 2023:2023.04.13.536711. doi:10.1101/2023.04.13.536711

70. Xia Y, Prokop S, Giasson BI. “Don’t Phos Over Tau”: recent developments in clinical biomarkers and therapies targeting tau phosphorylation in Alzheimer’s disease and other tauopathies. Molecular Neurodegeneration. 2021;16(1):37. doi:10.1186/s13024-021-00460-5

71. Hu W, Zhang X, Tung YC, Xie S, Liu F, Iqbal K. Hyperphosphorylation determines both the spread and the morphology of tau pathology. Alzheimers Dement. 2016;12(10):1066–1077. doi:10.1016/j.jalz.2016.01.014

72. Robert A, Schöll M, Vogels T. Tau Seeding Mouse Models with Patient Brain-Derived Aggregates. International Journal of Molecular Sciences. 2021;22(11):6132. doi:10.3390/ijms22116132

